# A framework to trace microbial engraftment at the strain level during fecal microbiota transplantation

**DOI:** 10.1101/2022.05.18.492592

**Authors:** Yiqi Jiang, Shuai Wang, Yanfei Wang, Xianglilan Zhang, Shuaicheng Li

**Affiliations:** Department of Computer Science, City University of Hong Kong, 83 Tat Chee Ave, Kowloon Tong, Hong Kong, China; State Key Laboratory of Pathogen and Biosecurity, Beijing Institute of Microbiology and Epidemiology, Beijing 100071, China

**Author notes:** Equal contributor (Yiqi Jiang, Shuai Wang).

**Keywords:** Fecal microbiota transplantation, Metagenomics, microbial strains

## Abstract

**Background:** Fecal microbiota transplantation (FMT) may treat microbiome-associated diseases effectively. However, the mechanism and pattern of the FMT process require expositions. Previous studies indicated the necessity to track the FMT process at the microbial strain level. At this moment, shotgun metagenomic sequencing enables us to study strain variations during the FMT.

**Result:** We implemented a software package PStrain-tracer to study microbial strain variations during FMT from the shotgun metagenomic sequencing data. The package visualizes the strain alteration and traces the microbial engraftment during the FMT process. We applied the package to two typical FMT datasets: one ulcerative colitis (UC) dataset and one *Clostridium difficile* infection (CDI) dataset. We observed that when the engrafted species has more than one strain in the source sample, 99.3% of the engrafted species will engraft only a subset of strains. We further confirmed that the all-or-nothing manner unsuited the engraftment of species with multiple strains by heterozygous single-nucleotide polymorphisms (SNPs) count, revealing that strains prefer to engraft independently. Furthermore, we discovered a primary determinant of strain engrafted success is their proportion in species, as the engrafted strains from the donor and the pre-FMT recipient with proportions 33.10 % (*p*-value = 6*e −* 06) and 37.08 % (*p*-value = 9*e −* 05) significantly higher than ungrafted strains on average, respectively. All the datasets indicated that the diversity of strains bursts after FMT and decreases to one after eight weeks for twelve species. Previous studies neglected strains with their corresponding species showing insignificant differences between different samples. With the package, from the UC dataset, we successfully determined the strain variations of the species *Roseburia intestinalis*, a beneficial species reducing intestinal inflammation, colonized in the cured UC patient being engrafted from the donor, even if the patient hosted the same species yet before treatment. We found seven strains in donors from the CDI dataset and one strain in pre-FMT recipients from eight species that associated CDI FMT failure.

**Conclusion:** PStrain-tracer is the first framework that tracks strain alterations in metagenomic sequencing data of FMT. PStrain-tracer implemented several methods specialized for FMT experiment samples, such as visualization of strains abundance alteration in the FMT experiment and determinant strains detection in FMT failure. We applied PStrain-tracer on two published datasets, uncovered novel strains related to FMT failure, and demonstrated the necessity of analyzing the whole-genome shotgun metagenomic data of FMT at the strain level. We also developed an online visualizer of PStrain-tracer for the users to adjust their visualized results online. The package is available at https://github.com/deepomicslab/PStrain-tracer.

## Background

Fecal microbiota transplantation (FMT) treatments transfer the fecal material from a healthy donor to the recipient with microbiome-associated diseases. They achieved successful instances to treat *Clostridium difficile* infection (CDI) [1−5] and ulcerative colitis (UC)[6−8]. Moreover, FMTs have therapeutic potential to treat other gut microbiome-associated diseases, including Crohn’s disease [9], irritable bowel syndrome [10], autism [11], autoimmune disorders [12], and metabolic syndrome [13, 14]. However, there still demands a deep understanding of the underlying mechanism and pattern in FMT treatments to enhance the usage or reduce the adverse outcomes of FMT [15]. Therefore, some efforts have been made to explore the mechanism by studying animal models [16].

Next-generation metagenomic sequencing provides us the opportunity to learn the microbial composition for each sample. Thus, we can track the microbial composition changes at a proper level of resolution. Using 16S ribosomal RNA genes amplicons sequencing could give rise to a better understanding of FMT [17]. Furthermore, researchers started to use shotgun metagenomic sequencing data to analyze FMT. Some studies mapped the sequencing reads to the reference to track the changes during the FMT process [18]. Other researchers tried to assemble the genomes to get direct access to the genomic content of each sample. However, they could only reconstruct metagenome-assembled genomes (MAGs) instead of the complete genomes [19]. It is challenging to assemble different highly similar strains for each species [20]. The taxonomy annotations of MAGs could not reach a high resolution. Most MAGs got the taxonomy annotations at order levels. [19]

Previous research showed that bacterial differences in the gut at the strain level of the same species could lead to human diseases or different physiological activities. Most *Escherichia coli* strains residents in the gut are harmless, while O157:H7 is an acute enterohemorrhagic pathogen, and pks+ shows long-term risks of colorectal cancer [21, 22]. A specific methicillin-resistant *Staphylococcus aureus* strain revealed an increased virulence[23]. Amandine *et al*. in mouse gut microbiome models found specific strains of *Akkermansia muciniphila* could control the metabolic conditions of diet-induced obesity[24]. Yaowen *et al*. detected *Bacteroides coprocola* colonized in type 2 diabetic patients had specific SNPs[25]. Therefore, exploring the FMT process at the strain level is necessary.

Strain Finder is a tool developed by Smillie *et al*., designed to infer strains and had been applied in a whole genome shotgun metagenomic dataset of FMT samples [26]. They found several patterns of strain engraftment between the donor and patients. For instance, they uncovered that strains of the same species engraft in an all-or-nothing manner. However, the reliability of these findings depends on the performance of the strain inference tool.

We previously developed PStrain [27] to improve the performance of strain inference. It infers strains based on the single-nucleotide variants (SNVs) on the MetaPhlAn2 marker genes [28]. It phases the variants using the genotype frequencies at each locus. Moreover, it introduces second-order genotype frequencies to adapt the information of reads which cover multiple loci. PStrain utilizes an iterative optimization algorithm that integrates linear programming and dynamic programming to infer strains.

We implemented a package to track microbial colonization at the strain level to understand what happened in the gut microbiome during the FMT experiment. The purpose of the package is to solve two commonly concerned points in FMT research: the potential FMT failure-related strains in the donor and the pre-FMT recipient, and the source of the engrafted strains in post-FMT recipient. Firstly, it incorporates the PStrain package to profile the strains’ sequences and abundance. The package could visualize the strain proportion alteration and trace the microbial engraftment in FMT experiments. Based on the provided patients’ relief situation after FMT treatment, the package could detect the potential strains involved in the effectiveness of FMT. Moreover, the package will report the most similar strains in the public genome database of these potential strains for the user to check the genome function further. We also developed an online visualizer using the output of PStrain-tracer. The visualizer enables the user to adjust the graph online, like sorting samples, filtering samples and recoloring.

We applied the package to a CDI FMT dataset with 74 samples and a dataset of UC FMT with ten samples. We detected eight strains associated with the CDI FMT failure, which had been neglected before as their corresponding species abundance has no difference between failure and success FMT experiment samples. We could distinguish the engrafted strains were sourced from the donor or the pre-FMT recipient with strain genotypes. Then we found that the colonization of *Bacteroidales* and *Clostridiales* might be vital to cure UC. Finally, we observed two general multiple strains related patterns in both datasets. Firstly, the strains of the same species do not always engraft in an all-or-nothing manner. The strains were more frequently partially engrafted in the post-FMT samples than completed engrafted. Besides, the engrafted strains have a significantly higher proportion than the rest in the same species. Secondly, the strain diversity would burst after FMT and then reduce to one strain in twelve species.

## Results

### Strains associated to FMT failures

We tried to determine if the strain-level resolution will reveal more causes of FMT failure in dataset CDI. This dataset contains 22 FMT experiments involving seven donors samples, 22 pre-FMT recipients samples, and 45 samples at different time points after transferring (post-FMT). The case treated had no diarrhea after eight weeks after FMT was referred to as an FMT success; otherwise, it was an FMT failure. We can divide the donor samples into two groups; the first group includes FMT02, FMT04, and FMT27, where they lead the six FMT failures, and the second group contains FMT01, FMT30, FMT33, and FMT34. Denote the two groups as donor-F and donor-S, respectively. Though donor-F samples lead to nine FMT successes, we supposed the microbial composition in donor-F samples may work as the potential determinant of FMT failure. The recipients were divided based on the FMT results into two groups recipient-F and recipient-S. We performed PStrain and our package on this dataset to infer strains and analyze the results.

To explore the results that species-level analyses could not afford, we reserved the strains fit: (1) species contains more than one strain; (2) the strain exists in at least two samples. We obtained 34 strains in all the donors after the filtering, and then we tried to find strains associated with FMT failures.

We visualized the donor samples with the abundance of the 34 strains; the result indicated the strains from the different groups (donor-F and donor-S) varies heavily (Figure 1A). It showed that strains from the 34 strains might be vital to the FMT treatment result.

**Figure 1.**
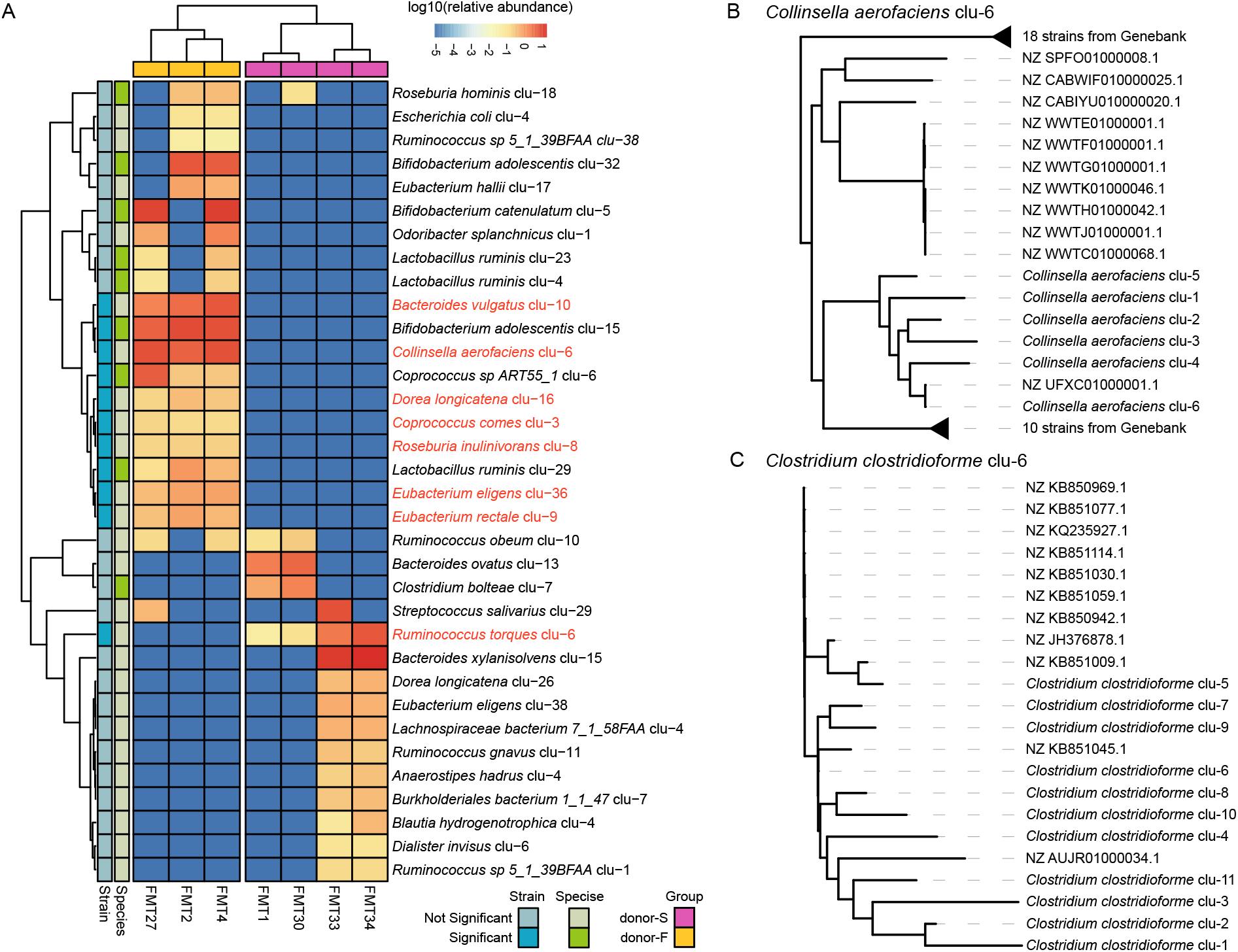
The existence of certain strains may cause FMT failure. (A) The heatmap of strain abundance in group donor-F and group donor-S. The annotation of strains shows whether they were significantly different between the two groups, and the abundance of their corresponding species was significantly different between groups or not. The strains whose abundance was significantly different while had no significant difference in the abundance of related species were highlighted in red. We constructed a phylogenetic tree of the strains we highlighted in (A), respectively. We used all strains we detected and all assemblies in GenBank of certain species. We could find a known assembly in the same branch for two strains, *Collinsella aerofaciens clu-6* (B) and *Clostridium clostridioforme clu-6* (C).

We detected 21 strains that show significantly different abundance between the two groups. We discarded the strains whose corresponding species’ abundance was significantly different between the two groups (Wilcox.test, *p <* 0.05). In the remaining eight strains, *Ruminococcus torques clu-6* was enriched in donor-S, the other seven strains were enriched in donor-F. *Bacteroides vulgatus* enriched in donor-F is a known putative pathogenic bacteria in IBD [29, 30].

In the comparison of recipient-F and recipient-S, only the abundance of strain *Clostridium clostridioforme clu-6* was significantly enriched in recipient-F (Wilcox.test, *p* = 0.022) while the abundance of the species was not significantly different.

Further, we determined if the strains with significantly different abundance were known strains or not. We downloaded all reported assemblies of certain species from GenBank, aligned the genomes to marker gene reference exacted the genotypes at SNV loci in all strains to construct the phylogenetic tree (Fasttree2, default setting). If the strain and an assembly were located on the same branch, we inferred that strain was close to the known strain. We found *Collinsella aerofaciens clu-6* close to *Collinsella aerofaciens* strain NCTC11838 (GenBank accession: NZ UFXC01000001.1) in donor-F and donor-S comparison. *Clostridium clostridioforme clu-6* was close to *Clostridium clostridioforme* strain 90A7’s genomic scaffold acsOa-supercont1.1 (GenBank accession: KB851045.1) in recipient-F and recipient-S comparison. Both of these two strains lack related clinical information.

We identified the strains that may impact the FMT results in both donors and recipients. To adopt FMT to cure CDI more reliably, we should learn the genome features of these strains in the future, such as gene contents, virulence factors, and antibiotic resistance genes, to investigate how these strains affect FMT.

### Tracking microbial colonization in FMT reveals strains related to intestinal inflammation remission

In dataset UC, two patients (R01 and R02) with ulcerative colitis got relief after FMT treatment. The dataset contains four samples for each experiment, the donor samples (DS01 and DS04), pre-FMT recipient samples (R01 pre-FMT and R02 pre-FMT), post-FMT samples collected four weeks after FMT (R01 W4 and R02 W4), and post-FMT samples collected eight weeks after FMT (R01 W8 and R02 W8). The bacteria in post-FMT samples had three possible sources: recipient, donor, or environment. The bacteria sourced from healthy donors colonized in patients after FMT may decrease intestinal inflammation. Therefore we tried to infer the strains sourced from donors and colonized in patients using PStrain and our package.

Some species were merely colonized in the donor and post-FMT samples. People may infer that they are transferred from donors to the patients. However, our results revealed that the inference could be imprecise. For instance, although *Paraprevotella clara* of R01 W4 only existed in the donor and post-FMT samples. The strains from the donor and post-FMT sample were different (Figure 2A). The species colonized in the post-FMT sample unlikely engrafted from the donor. This case suggested the necessity to infer engraftment of bacteria at the strain level.

**Figure 2.**
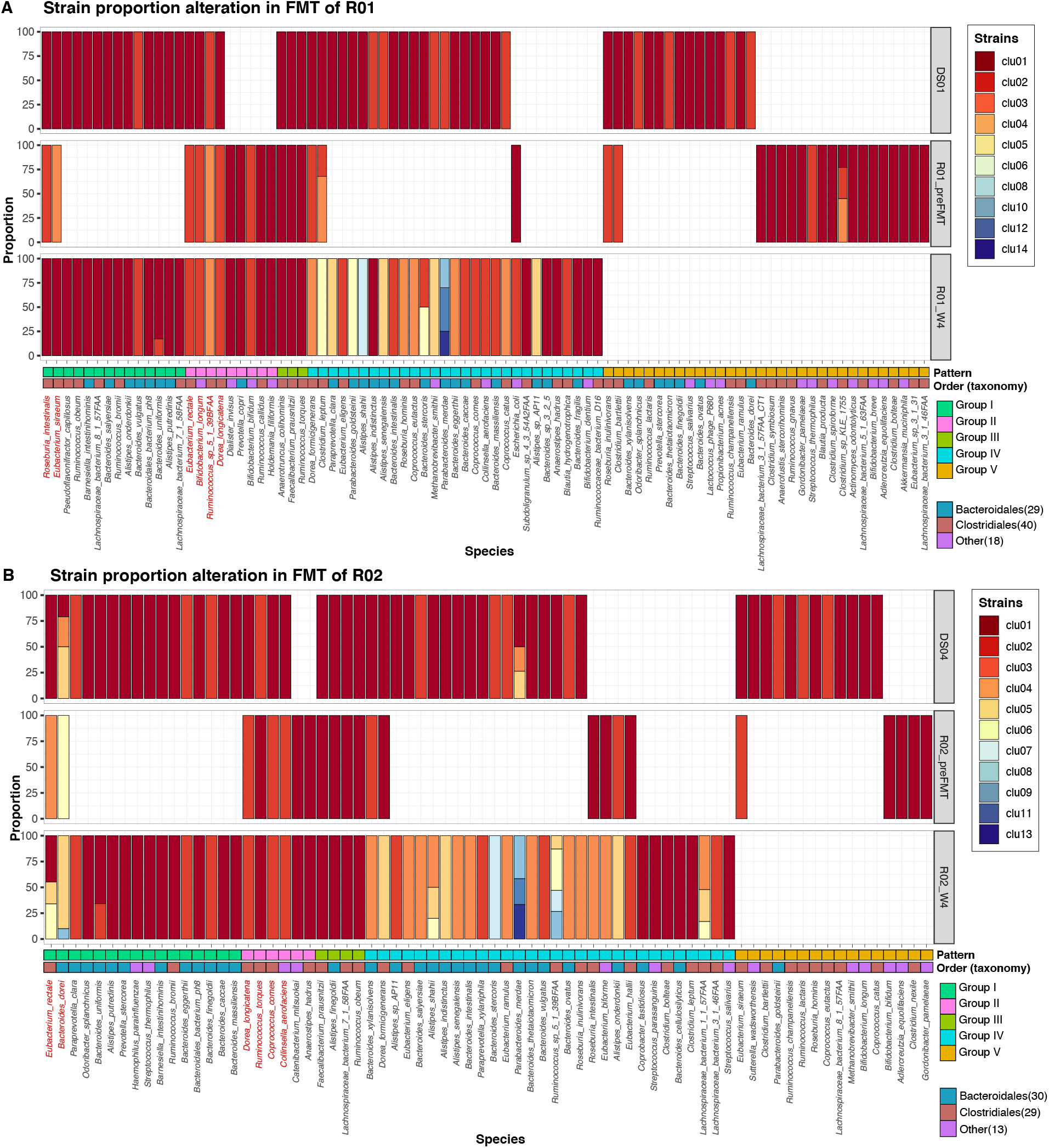
Strain proportion alteration in UC FMT process. The strain proportion alteration from donor and recipient to post-FMT samples in *R*01(A) and *R*02(B). The bar plot represents the strain proportion of each species. The different color of bar plot reflects different strains. Annotation bars of species show the engraftment pattern group and the taxonomy order. The red species name means those species existed in the donor, pre-FMT and post-FMT samples. However, we still could determine the source of strains in post-FMT based on our strain-level analysis.

We classified the strains that appear in the FMT experiment into five groups based on the engraftment pattern. If the strain in post-FMT was different from the strains in both pre-FMT and donor, we could not determine this species’ source (Group IV). Another case was when the donor, pre-FMT, and post-FMT samples contained the same strain, while we could not distinguish the strain source (Group III). Sometimes, the same species showed up in donor, recipient, and post-FMT samples, while the strains differed. We could distinguish the species engrafted from the donor to the patient if the strains colonized in donor and post-FMT are the same, while the strains in donor and recipient are different (Group I). In the same way, we could determine the strains sourced from the recipient (Group II). Also, we could notice several strains perished or colonized unsuccessfully in the recipient (Group V). Therefore, according to the strain-level results, we could profile the bacteria’s engraftment more precisely during the FMT.

We identified 59 and 69 strains in R01 W4 and R02 W4, respectively. Among them, 15 (25%) and 20 (29%) strains could be determined sourced from the donor (Group I, Figure 2) to post-FMT samples. The taxonomy of these strains was majorly the order *Bacteroidales* and *Clostridiales* both in R01 and R02 array. In the species that existed in the donor, pre-FMT and post-FMT samples jointly, we determined engrafted strains of *Roseburia intestinalis* and *Eubacterium siraeum* were sourced from the donor in the R01 FMT as the strains in post-FMT were the same as the donor and different from the recipient’s strains. Similarly, we could determine *Eubacterium rectale* and *Bacteroides dorei* were sourced from the donor in R02 FMT.

*R. intestinalis* is a known species related to inflammation in colitis. The flagellin derived by *R. intestinalis* is a negative regulator of intestinal inflammation. *R. intestinalis* inhibits interleukin-17 excretion and promotes regulatory T cells differentiation in colitis. Besides, the target colonization of *R. intestinalis* is a new colitis therapy strategy [31−34].

### The engrafted strains had significantly higher proportions than ungrafted strains

Unlike previous research Smillie *et al*. published, we observed that the engraftment of strains in the species with multiple strains does not always follow an all-or-nothing manner. When we consider all species from the donor or the recipient, the strains’ engraftment followed the all-or-nothing pattern. (Figure 3A-B). However, after we removed the species containing one strain, the partial engraftment is more frequent than all-or-nothing (Figure 3C-D). 24 of 25 donor engrafted species and 18 of 20 pre-FMT engrafted species were partial engraftment. Summary all engrafted species, we determined 93.3% species engrafted partially. Smillie *et al*. believed the all-or-nothing manner represents the strains in one species prefer transfer together as a cohesive unit. While our result revealed, the strains frequently engraft independently instead.

**Figure 3.**
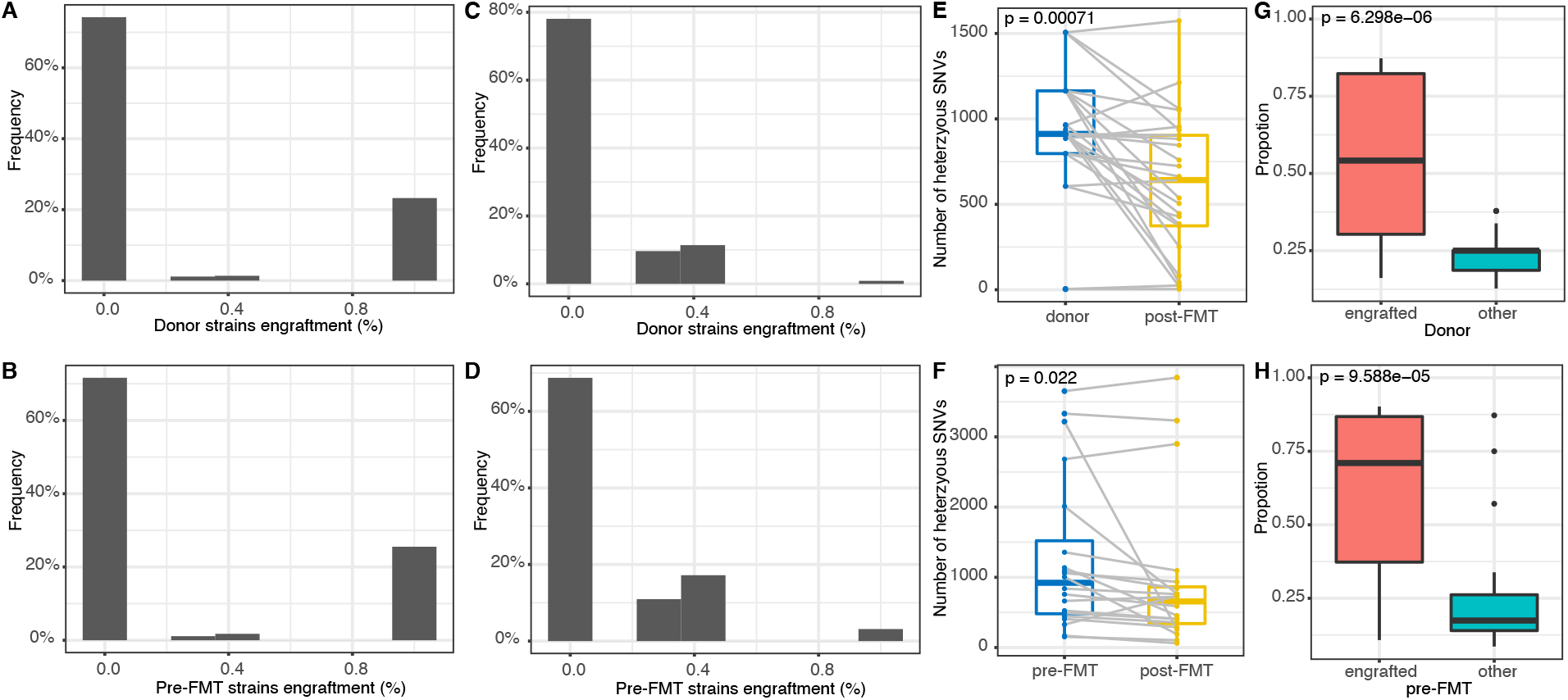
Most species engrafted partial strains from the donor to the first post-FMT samples. The proportion of the engrafted strains were significantly higher than the other strains. (A-B) The frequency of the percentages of strains from the donor and the pre-FMT were engrafted in the post-FMT samples. (C-D) After filtering the species contains one strain, redraw the frequency same as (A-B). (E-F) The heterozygous SNPs number alteration in the paired source and the first post-FMT sample. The grey lines link the paired samples, and the number increased in post-FMT samples was highlighted in red links. (G-H) Proportion comparison between engrafted strains and the other strains in the engrafted species. (A, C, E, G) are based on donor engrafted species (B, D, F, H) are based on pre-FMT engrafted species.

We confirmed partial engraftment with the number of heterozygous SNPs in the donor and the post-FMT samples. Besides *Ruminococcus torques* in FMT1 to FMT5 which the engrafted species in ‘all’ pattern, *os-eburia inulinivorans* in FMT2 to FMT21 and *Dorea longicatena* in FMT27 to FMT82, *Bifidobacterium adolescentis* in FMT2 to FMT15, and *Ruminococcus sp 5 1 39BFAA* and *Ruminococcus torques* in FMT1 to FMT35 were the only five had more heterozygous SNPs in the donor than in the post-FMT sample. These species had other strains with undetermined sources in the post-FMT samples. The rest 19 cases showed fewer heterozygous SNPs in post-FMT samples, which supported the number of strains decreasing and partially engraftment of these species. (Figure 3E) The number of heterozygous SNPs in recipient engrafted species decreased in 16 of 20 engrafted species in the post-FMT samples.(Figure 3F) The exceptions contained all engrafted species, and the species invoked new strains from the environment. The partial engraftment was more frequent than completed engraftment.

Since we have determined the independent colonization of strains, we further explored the characteristics of the engrafted strains, and the results showed these strains have significantly higher proportions than ungrafted strains. We inferred that 24 donor engrafted species and 18 pre-FMT recipient engrafted species partly transferred the strains to the post-FMT samples. We explored the difference between the engrafted strains and the rest in donors and pre-FMT recipients. Firstly, we compared the abundance of engrafted strains and the rest strains. We found there was no significant difference (Wlicox.test, *p* = 0.24 for recipients and *p* = 0.33 for donors). Furthermore, we counted and compared the proportions of engrafted strains and the rest strains of the same species.

The results indicated that the proportions of engrafted strains were significantly higher than the rest strains in the same species (Wlicox.test, both *p <* 0.001)(Figure 3G-H), which might suggest that the strains with a high proportion were more likely successfully transferred. While the determinant of a bacterial transmission was based on its species traits, as Smillie *et al*. mentioned [26].

### A brief burst of the strains diversity after FMT

After analyzing the strain result using our package, we found an intuitive pattern in both datasets. For some species, the strains’ diversity would increase after FMT and then decrease quickly.

In dataset UC, strain proportion alteration revealed the number of strains in several species raised at four weeks after FMT and stable to single strain at the point of 8 weeks after FMT. *Parabacteroides merdae* and *Bacteroides stercoris* in R01 array, *Ruminococcus sp 5 1 39BFAA, Alistipes shahii, Bacteroides dorei* and *Eubacterium rectale* in R01 array had more than one strains at W4 point, and single strain at W8 point (Figure 4C-D). Besides *Bacteroides dorei*, other species had only one strain in donor and pre-FMT recipient.

**Figure 4.**
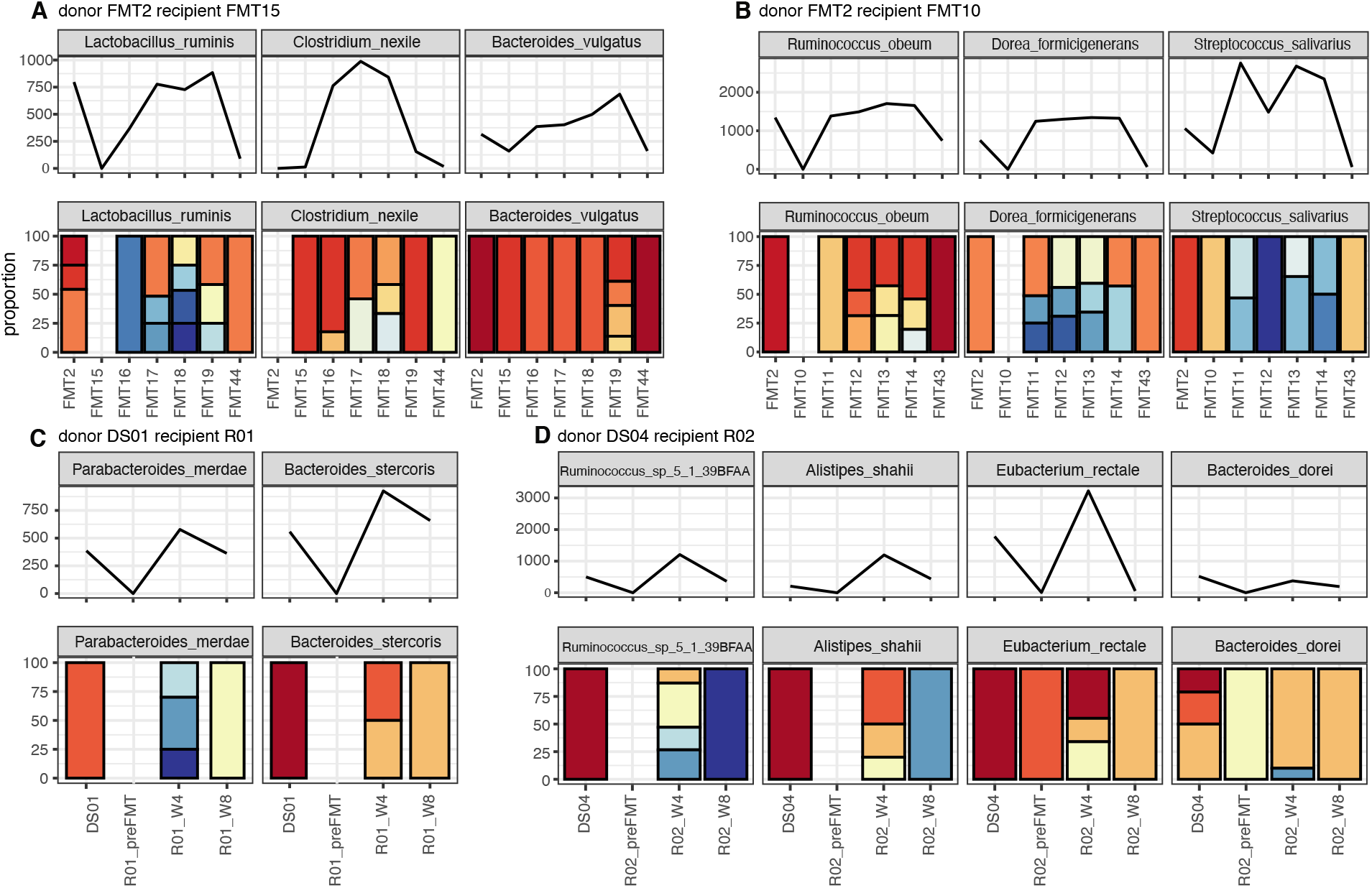
The diversity of strains will burst after FMT and finally stable to one. The strain proportion of species that had the number of strains raised after FMT and the number of strains became one in the last post-FMT samples. The color in proportions represented different strains. We displayed all samples in FMT experiments: the donor, the recipient, and all post-FMT samples. The number of heterozygous SNPs in corresponding samples was showed above the proportion bar. We showed FMT experiments of FMT15(A) and FMT10(B) in dataset CDI, R01(C), and R02(D) in dataset UC. The listed samples in each FMT experiment follow the order: the donor sample, the pre-FMT sample, the sorted posFMT samples based on the sampling days after FMT.

In dataset CDI, we discovered the same pattern: the number of strains increased in the early stage of post-FMT and stable to one strain later. Although the sampling time of post-FMT samples differs from which in dataset UC, 30 of 45 post-FMT samples were sampling two weeks after FMT. We still could uncover such conditions in this dataset. For instance, we showed this pattern in two FMT treatment experiments sampling at different processes, both of FMT experiments including one donor sample and five post-FMT samples.

In the FMT experiment that transplanted donor sample FMT2 to the recipient whose pre-FMT sample corresponded to FMT10, there were five post-FMT samples with the sampling day after FMT in brackets: FMT11 (day10), FMT12 (day10), FMT13 (day18), FMT14 (day18) and FMT43 (day83). We discovered *Dorea formicigenerans, Ruminococcus obeum*, and *Streptococcus salivarius* all had multiple strains in FMT11, FMT12, FMT13, and FMT14. (Figure 4B) However, the strain stable to a single strain again at the final post-FMT FMT43, which sampled 83 days after the transfer. In the FMT experiment that transplanted donor sample FMT2 to the recipient whose pre-FMT sample corresponded to FMT15, there were five post-FMT samples with the sampling day after FMT in brackets: FMT16 (day3), FMT17 (day3), FMT18 (day6), FMT19 (day6) and FMT44 (day90). *Ruminococcus obeum, Dorea formicigenerans* and *Streptococcus salivarius* showed the same pattern (Figure 4A).

We checked the heterozygous SNPs loci for these species. The number of heterozygous SNPs increased after FMT and decreased then, which consistently supported the pattern. The previous work reported the infection or colonization of *C*.*difficile* would reduce the diversity in the human gut microbiome [35]. The diversity increasing after FMT is consistent with previous studies [36−38]. We were the first time to report which strains play roles in the diversity burst after FMT. The brief burst of strains’ diversity might bring unknown risks to the patients. It is vital to explore the mechanism.

### Online visualization of the data

We developed an online visualizer FMT overview for the outputs of Pstrain-tracer. We used Oviz[39], a data visualization framework that fully utilizes the modern web browser, to deliver high-quality vector charts with smooth user interaction. We adopted Vue.JS to implement a functional editor for users to upload data and adjust chart settings. Accepting a strain proportion file from PStrain-tracer, and a metadata file of species. The visualizer would draw all samples’ strain proportion bar plots in rows and a metadata heatmap at the bottom. A tooltip containing the species, strain, and proportion will show when users hover on the bar. Users can select metadata as sorting criteria in the editor to reorder the species. The editor also provides functions such as filtering species and recoloring. We used the input files of Figure 2B as a demo. The online visualizer with a user manual is freely available on the GutMeta website (https://gutmeta.deepomics.org/visualizer/analysis/fmt-overview), running on a server owned by the City University Of Hong Kong.

## Discussion

### Comparison with MAG approach

The previous research on the dataset UC used the metagenome-assembled genomes (MAG) approach to track microbial colonization in FMT. The MAGs applied in the analysis were assembled from four samples of the only donor individual at different time points. The tracking of the MAGs colonization situation in post-FMT, while the taxonomy annotation of MAGs was poor. Only 31 of 93 MAGs got annotation at the genus level. Our approach is based on the MetaPhlAn2 marker gene database. We could analyze the data and track the detailed changes at the strain level. The strain-level approach makes it possible to distinguish whether the strains in post-FMT were sourced from donor or recipient, even if both donor and recipient samples contained a particular species.

We successfully inferred seven species’ engraftment while the donor, recipient, and post-FMT samples all had the same species. Six of seven species were sourced from the recipient, while one species, *Roseburia intestinalis*, was sourced from the donor in the *R*01 array.

### Reject the engraftment pattern of all-or-nothing

We analyzed the strain’s engraftment pattern in the species that comprised at least two strains in the donor or pre-FMT recipient. Our results rejected previous all-or-nothing engrafted conclusions in two aspects. Firstly, we counted the number of heterozygous SNPs for the engrafted species and found the number of heterozygous SNPs decreased after FMT. If the species engrafts following an all-or-nothing pattern, the number of heterozygous SNPs will keep stable during the FMT. However, if the species engrafts partial strains, the number of heterozygous SNPs will decrease. The results supported the second case; therefore, we could conclude that the species does not engraft in an all-or-nothing manner. Secondly, we inferred strains using PStrain, and the results directly indicated that most of the species engraft partial strains to post-FMT samples. Besides, we uncovered a pattern that the engrafted strains have higher proportions than the remaining strains in the same species. Therefore, we concluded that the engrafted species with multiple strains not following the all-or-nothing manner.

Also, we analyzed the reason that Smillie *et al*. observed the all-or-nothing pattern. Smillie *et al*. designed their strain-inferring tool following an intuition that SNPs caused by the same strain in different samples will show roughly equivalent frequencies. Intuitively, this assumption will lead to similar strain frequencies across samples. Based on these results, one may get a false conclusion while looking into the similarity of strain frequencies between the donor and post-FMT samples. However, PStrain profiles strain with no assumption of SNPs or strain frequencies across samples. Therefore, it is more reliable to study microbial engraftment across samples based on PStrain’s results.

### Limitation and future work

The major disadvantage of our method is the fixed threshold in clustering strains. In 16S rRNA-based analysis, identity lower than 97% was considered as different species[40]. However, we used the marker genes with a lower conservative than the 16S rRNA gene. Currently, there are no accepted standards of identity strain from the marker genes. First, different bacteria usually had different mutation rates, and species specialized strains identification threshold will help strain clustering. Secondly, we used a marker genes database. The length of the marker gene occupied different portions of the total length of the genome in different species. In the future, we intend to build a math model to evaluate the strains clustering threshold in different species.

Moreover, we should consider preserving the heritage relationship between strains in the evaluation process. For instance, if the mutation ratio exceeds the strain clustering threshold, the donor’s strain will be considered a new strain and can not be inferred source. Therefore, we could try to get an optimal threshold in clustering for a particular species to reserve more heritage relationships. Alternatively, we could develop a method to infer more heritage relationships in the tracing stage after strain clustering. It would reduce the proportion of unknown sourced strains in the post-FMT samples.

Another factor we should consider in the further analysis is the definition of FMT failure. For the CDI dataset we used, the original data publishers had defined the FMT success as no relapse of diarrhea after eight weeks from FMT. However, other works did not have such an explicit time limitation [41−44]. If we collected FMT failure samples from other projects and merged them, the detected determinant strains would be affected. The precise and clear definition of FMT failure needs to be discussed and determined by professional medical researchers in the future. From the perspective of bioinformatics analysis, we could consider using remove batch effects methods or build a classifier to filter the samples that participated in the comparison.

## Conclusions

We present a framework to analyze FMT at the strain level. With the framework, we could present the strain alterations in every FMT experiment, determine the source of engraftment strains in the post-FMT samples and discover the potential determinant strains of FMT failure. We summarize the scripts used in the framework to a package, PStrain-tracer. We applied the package in two published FMT datasets, and we found eight strains possibly related to CDI FMT failure; the engrafted strains usually have higher proportions than the rest of the strains, and the strains diversity has a brief burst after FMT. Finally, we developed a web visualizer to present the result from PStrain-tracer online.

## Methods

### The framework to trace FMT at the strain level

The package took the whole-genome shotgun metagenomic sequencing data of *fastq* format of a sample as input. First, PStrain [27] inferred the strain genotypes and abundance of each species in each sample. Then we cluster the strains obtained in all the samples. After that, we used the cluster name to represent the corresponding strains and calculate the cluster’s abundance by summing up the corresponding strains’ abundance in each sample. Finally, we outputted the strain proportions in all samples and visualized the strain proportion alteration for every FMT experiment, respectively (Figure 5A).

**Figure 5.**
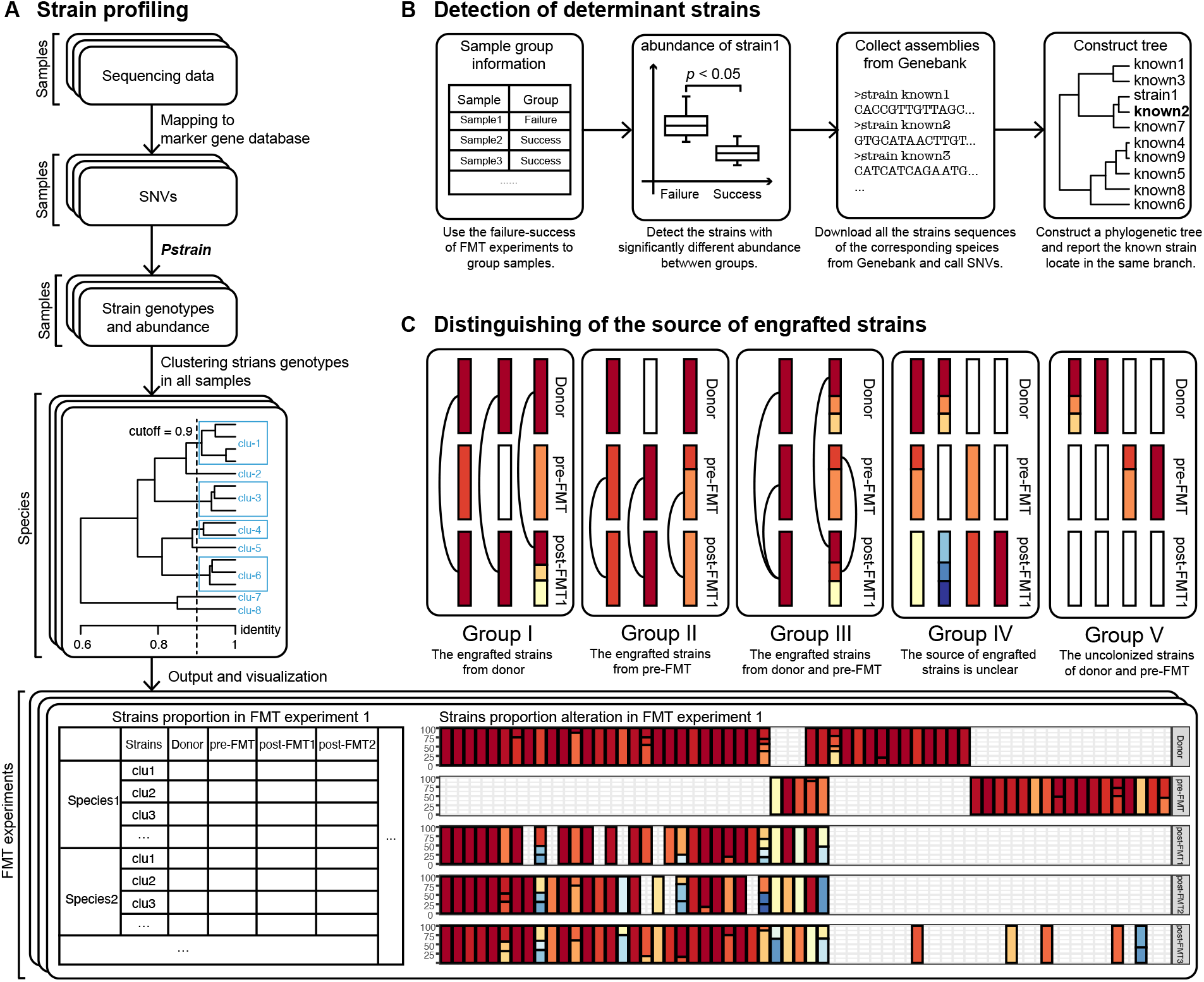
The flowchart of our FMT strain-level analysis package. (A) Profile strains for the samples using PStrain. Map the reads to the MetaPhlAn2 marker genes, call SNV, and infer strains genotypes and abundance. Then cluster the strains obtained in all samples and calculate the abundance of strain clusters in each sample. (B) Infer differential strain between FMT success and failure. Divide the samples into groups, compare the strains’ abundance between groups to explore vital strains, and find the close assembly sequence from the GenBank. (C) Infer bacterial engraftment during the FMT. The strains that showed up in the FMT process are divided into five groups. We infer the engraftment of species groups based on the pattern.

We adopted several steps to explore the determinant strains between groups: first, we assigned the samples into groups according to the phenotypes. In our approach, the groups were FMT success and FMT failure, as we concern most. Then, we assumed the strains significantly different between groups might be phenotype-associated and identify these strains. Subsequently, we collected all the assemblies of corresponding species from GeneBank and selected the closest assembly sequence for the concerned strains (Figure 5B). We could infer the source of engrafted strains based on genotypes, and we grouped species into five groups based on the strain existence in the donor, pre-FMT, and post-FMT samples (Figure 5C) to present the engraftment pattern. We also provided the functions such as tracing the alteration of the number of strains, and proportion comparison of strains, to evaluate the characteristic in strains engraftment.

We will introduce the methods in detail in the subsequent sections. The scripts of every analysis are contained in our package.

### Profiling strains with PStrain

Based on the PStrain [27] preprocessing pipeline (Figure 5A), the sequencing data were first aligned to the MetaPhlAn2 marker gene database (version mpa v20) to obtain the species abundance in each sample [28]. We performed PStrain with default parameters for each sample to infer strains for species with a depth larger than 5x and otherwise, we inferred species’ consensus sequence. After that, we cluster the strains inferred from all samples in both datasets with hierarchical clustering. We clustered strains based on the genotypes at all SNPs loci with the similarity cutoff of 90%. The proportion of the cluster is the sum of the proportion of strains in the cluster. We denoted the strains in each sample as their corresponding cluster name.

### Detect the strains associated to FMT failure

#### Detect the strains that differ between FMT success and failure

To explore the strain difference between samples of FMT success and failure, we divided the donors into two groups: donor-S and donor-F (Figure 5B). The donors in donor-S successfully helped the recipients cure the CDI by transferring their microbiome. In contrast, donors in donor-F failed to help cure recipients’ CDI. Similarly, we also divided the pre-FMT recipient samples into two groups: recipient-S and recipient-F. The difference between the groups may be vital to the FMT treatment results.

We removed the species that meet either one of the following two conditions: the species merely existed in a single sample or only consisted of one strain in all samples. We plotted the heatmap for the samples with strains’ proportion by pheatmap package (version 1.0.12) of R (version 3.6.1) [45, 46]. For each strain, we compared the strains’ abundance and the abundance of corresponding species between groups using the Wilcox.test by R. We chose the strains with significantly different (*p−value <* 0.05) abundance and nonsignificantly different abundance of its corresponding species between groups as the strains associated with FMT failure.

#### Reporting close assembly from GenBank for the strains

For the strains associated with FMT failure, which we determined above, we downloaded all accessible assembly sequences of its corresponding species from Genebank [47]. We extracted the marker gene sequences of the species in the MetaPhlAn2 database as the reference sequence [28]. We detected the homology regions and the SNVs on these regions in each assembly by aligning the assembly sequence to the reference sequence with MUMmer (version 3.20) [48]. The pipeline was: “nucmer −mum” for alignment, “delta-filter -1” for filtering homology regions, “show-snps - CIrT” to extract SNVs in homology regions. Then, we used in-house-built scripts to extract genotypes at all SNV loci from multiple sequence alignment results. Finally, we used an approximately maximum likelihood method of Fasttree2 (version 2.1.10) [49] to construct the evolution tree and report the most similar strain of a specific strain.

### Analysis of strains engraftment

#### Inference of strains engraftment during FMT

The strains colonized in post-FMT samples are from three sources: donor, pre-FMT recipient, or environment. However, we could not detect some strains due to the low abundance in the donor or pre-FMT recipient. Therefore, we merely focused on the detected strains currently. We could clarify the source of engrafted strains (Figure 5C), even both donor and pre-FMT recipient samples contained the particular species. If the strain genotypes in the post-FMT samples were the same as the strain genotypes in the donor, we could determine the strain sourced from the donor (Group I). In the same way, we could determine the strains sourced from the recipient (Group II). The strains were labeled with the uncertain resource in two ways: donor, pre-FMT, and the post-FMT samples contained the same strain (Group III); donor, pre-FMT, and post-FMT samples contained different strains (Group IV). The novel strains in post-FMT samples could come from the environment. Alternatively, they were undetected in the donor or pre-FMT recipient due to the low abundance. The strains mutated frequently beyond the similarity cutoff we set was the other possible explanation of Group IV. Also, we could notice several strains colonized in the pre-FMT recipient or donor not transferred to post-FMT (Group V).

#### Proportion difference between engrafted and ungrafted strains

The strain’s proportion was the ratio of the specific strain’s abundance in the sum of all strains’ abundance for the same species. We collected the proportions of strains from the results generated by PStrain. We focused on the engrafted species with multiple strains in the donor or pre-FMT recipient. For each species, we collected the proportions of engrafted and ungrafted strains, respectively. Then we plotted the box plot MAG metagenome-assembled genomes with ggplot2 package (version 3.3.0) [50] in R, and per-NCBI National Center for Biotechnology Information formed the differential analysis with Wilcox.test by R.

#### Tracing the alteration of the number of strains

We visualized the number of strains alteration in the donor, recipient, and post-FMT samples. We observed the number of strains, that is, the diversity of the corresponding species burst after FMT. We further counted and visualized the heterozygous SNVs loci in the samples. In the package, we provided a function to visualize the simultaneous changes of the number of strains and heterozygous SNVs loci.

### Data

The first FMT dataset we used was UC, which contains ten samples from two UC FMT experiments [19]. Dataset UC comprised four donor samples (DS01, DS02, DS03, DS04) from the same donor and three samples from each of the two recipients (R01, R02). Recipient samples were collected at the points pre-FMT, four weeks after FMT, and eight weeks after FMT. Both recipients had mild/moderate UC (R01 had proctitis, and R02 had left-sided/distal colitis), and both showed a decrease in intestinal inflammation after FMT. DS01 was transferred to R01, and DS04 was transferred to R02, while the donor samples DS02 and DS03 did not participate in FMT experiments. We noted samples involved in R01 or R02 FMT experiment as corresponding recipients array. The R01 array contains DS01, R01 pre-FMT, R01 W4, and R01 W8 and the R02 array contains DS04, R02 pre-FMT, R02 W8, and R02 W8.

Dataset CDI comprised samples related to FMT success and failure [26]. The FMT success was defined as no relapse of diarrhea after eight weeks from FMT in their work. Otherwise, it was an FMT failure. This dataset contains 74 samples of 22 FMT experiments: 22 recipients samples, seven donor samples, and 45 post-FMT samples. The seven donor samples were from 4 donors. The 22 recipient samples of 19 patients with CDI were recruited in the project, as three of them redid FMT after failure. 22 FMT experiments consist of 6 failure FMT and 16 success FMT. The sampling time of post-FMT samples was at specific time points ranging from 0 to 135 days after FMT. We downloaded the sequencing data of the dataset UC and CDI from the NCBI SRA database with the accession number PRJNA353655 and PR-JEB23524 (https://www.ncbi.nlm.nih.gov/sra).

#### Abbreviations

FMT: fecal microbiota transplantation
UC: ulcerative colitis
IBD: inflammatory bowel disease
CDI: *Clostridium difficile* infection
MAG: metagenome-assembled genomes
NCBI: National Center for Biotechnology Information

## Declarations

Ethics approval and consent to participate

Not applicable.

### Consent to publish

Not applicable.

### Availability of data and material

Datasets UC and CDI were downloaded from the NCBI SRA database with the accession number of PRJNA353655 and PRJEB23524. The package is available at https://github.com/deepomicslab/PStrain-tracer.

### Author’s contributions

SL designed the research. YJ and SW performed the experiments and wrote the manuscript. YW made the online visualizer and wrote the corresponding section in result. SL, YJ, SW, YW, and XZ reviewed the manuscript. All authors read and approved the final manuscript.

## Acknowledgements

We would like to thank Xinyao Miao who provides us the valuable support during the research.

## References

[1] Rupnik, M., Wilcox, M.H., Gerding, D.N.: Clostridium difficile infection: new developments in epidemiology and pathogenesis. Nature Reviews Microbiology 7(7), 526−536 (2009)

[2] Rohlke, F., Surawicz, C.M., Stollman, N.: Fecal flora reconstitution for recurrent clostridium difficile infection: results and methodology. Journal of clinical gastroenterology 44(8), 567−570 (2010)

[3] Gough, E., Shaikh, H., Manges, A.R.: Systematic review of intestinal microbiota transplantation (fecal bacteriotherapy) for recurrent clostridium difficile infection. Clinical infectious diseases 53(10), 994−1002 (2011)

[4] Van Nood, E., Vrieze, A., Nieuwdorp, M., Fuentes, S., Zoetendal, E.G., de Vos, W.M., Visser, C.E., Kuijper, E.J., Bartelsman, J.F., Tijssen, J.G., et al.: Duodenal infusion of donor feces for recurrent clostridium difficile. New England Journal of Medicine 368(5), 407−415 (2013)

[5] Lübbert, C., John, E., von Müller, L.: Clostridium difficile infection: guideline-based diagnosis and treatment. Deutsches Ärzteblatt International 111(43), 723 (2014)

[6] Anderson, J., Edney, R., Whelan, K.: Systematic review: faecal microbiota transplantation in the management of inflammatory bowel disease. Alimentary pharmacology & therapeutics 36(6), 503−516 (2012)

[7] Kunde, S., Pham, A., Bonczyk, S., Crumb, T., Duba, M., Conrad Jr, H., Cloney, D., Kugathasan, S.: Safety, tolerability, and clinical response after fecal transplantation in children and young adults with ulcerative colitis. Journal of pediatric gastroenterology and nutrition 56(6), 597−601 (2013)

[8] Moayyedi, P., Surette, M.G., Kim, P.T., Libertucci, J., Wolfe, M., Onischi, C., Armstrong, D., Marshall, J.K., Kassam, Z., Reinisch, W., et al.: Fecal microbiota transplantation induces remission in patients with active ulcerative colitis in a randomized controlled trial. Gastroenterology 149(1), 102−109 (2015)

[9] Zhang, F., Luo, W., Shi, Y., Fan, Z., Ji, G.: Should we standardize the 1,700-year-old fecal microbiota transplantation? American Journal of Gastroenterology 107(11), 1755 (2012)

[10] Borody, T.J., George, L., Andrews, P., Brandl, S., Noonan, S., Cole, P., Hyland, L., Morgan, A., Maysey, J., Moore-Jones, D.: Bowel-flora alteration: a potential cure for inflammatory bowel disease and irritable bowel syndrome? Medical Journal of Australia 150(10), 604−604 (1989)

[11] Hsiao, E.Y., McBride, S.W., Hsien, S., Sharon, G., Hyde, E.R., McCue, T., Codelli, J.A., Chow, J., Reisman, S.E., Petrosino, J.F., et al.: Microbiota modulate behavioral and physiological abnormalities associated with neurodevelopmental disorders. Cell 155(7), 1451−1463 (2013)

[12] Çorapçioğlu, D., Tonyukuk, V., Kiyan, M., Yilmaz, A.E., Emral, R., Kamel, N., Erdoğan, G.: Relationship between thyroid autoimmunity and yersinia enterocolitica antibodies. Thyroid 12(7), 613−617 (2002)

[13] Turnbaugh, P.J., Hamady, M., Yatsunenko, T., Cantarel, B.L., Duncan, A., Ley, R.E., Sogin, M.L., Jones, W.J., Roe, B.A., Affourtit, J.P., et al.: A core gut microbiome in obese and lean twins. nature 457(7228), 480−484 (2009)

[14] Vrieze, A., Van Nood, E., Holleman, F., Salojärvi, J., Kootte, R.S., Bartelsman, J.F., Dallinga−Thie, G.M., Ackermans, M.T., Serlie, M.J., Oozeer, R., et al.: Transfer of intestinal microbiota from lean donors increases insulin sensitivity in individuals with metabolic syndrome. Gastroenterology 143(4), 913−916 (2012)

[15] Chehoud, C., Dryga, A., Hwang, Y., Nagy-Szakal, D., Hollister, E.B., Luna, R.A., Versalovic, J., Kellermayer, R., Bushman, F.D.: Transfer of viral communities between human individuals during fecal microbiota transplantation. MBio 7(2) (2016)

[16] De Palma, G., Lynch, M.D., Lu, J., Dang, V.T., Deng, Y., Jury, J., Umeh, G., Miranda, P.M., Pastor, M.P., Sidani, S., et al.: Transplantation of fecal microbiota from patients with irritable bowel syndrome alters gut function and behavior in recipient mice. Science translational medicine 9(379) (2017)

[17] Angelberger, S., Reinisch, W., Makristathis, A., Lichtenberger, C., Dejaco, C., Papay, P., Novacek, G., Trauner, M., Loy, A., Berry, D.: Temporal bacterial community dynamics vary among ulcerative colitis patients after fecal microbiota transplantation. American Journal of Gastroenterology 108(10), 1620−1630 (2013)

[18] Li, S.S., Zhu, A., Benes, V., Costea, P.I., Hercog, R., Hildebrand, F., Huerta-Cepas, J., Nieuwdorp, M., Salojärvi, J., Voigt, A.Y., et al.: Durable coexistence of donor and recipient strains after fecal microbiota transplantation. Science 352(6285), 586−589 (2016)

[19] Lee, S.T., Kahn, S.A., Delmont, T.O., Shaiber, A., Esen, Ö.C., Hubert, N.A., Morrison, H.G., Antonopoulos, D.A., Rubin, D.T., Eren, A.M.: Tracking microbial colonization in fecal microbiota transplantation experiments via genome-resolved metagenomics. Microbiome 5(1), 50 (2017)

[20] Nielsen, H.B., Almeida, M., Juncker, A.S., Rasmussen, S., Li, J., Sunagawa, S., Plichta, D.R., Gautier, L., Pedersen, A.G., Le Chatelier, E., et al.: Identification and assembly of genomes and genetic elements in complex metagenomic samples without using reference genomes. Nature biotechnology 32(8), 822−828 (2014)

[21] Figler, H.M., Dudley, E.G.: The interplay of escherichia coli o157: H7 and commensal e. coli: the importance of strain-level identification. Expert review of gastroenterology & hepatology 10(4), 415−417 (2016)

[22] Cuevas-Ramos, G., Petit, C.R., Marcq, I., Boury, M., Oswald, E., Nougayréde, J.-P.: Escherichia coli induces dna damage in vivo and triggers genomic instability in mammalian cells. Proceedings of the National Academy of Sciences 107(25), 11537−11542 (2010)

[23] Gill, S.R., Fouts, D.E., Archer, G.L., Mongodin, E.F., DeBoy, R.T., Ravel, J., Paulsen, I.T., Kolonay, J.F., Brinkac, L., Beanan, M., et al.: Insights on evolution of virulence and resistance from the complete genome analysis of an early methicillin-resistant staphylococcus aureus strain and a biofilm-producing methicillin-resistant staphylococcus epidermidis strain. Journal of bacteriology 187(7), 2426−2438 (2005)

[24] Everard, A., Belzer, C., Geurts, L., Ouwerkerk, J.P., Druart, C., Bindels, L.B., Guiot, Y., Derrien, M., Muccioli, G.G., Delzenne, N.M., et al.: Cross-talk between akkermansia muciniphila and intestinal epithelium controls diet-induced obesity. Proceedings of the national academy of sciences 110(22), 9066−9071 (2013)

[25] Chen, Y., Li, Z., Hu, S., Zhang, J., Wu, J., Shao, N., Bo, X., Ni, M., Ying, X.: Gut metagenomes of type 2 diabetic patients have characteristic single-nucleotide polymorphism distribution in bacteroides coprocola. Microbiome 5(1), 1−7 (2017)

[26] Smillie, C.S., Sauk, J., Gevers, D., Friedman, J., Sung, J., Youngster, I., Hohmann, E.L., Staley, C., Khoruts, A., Sadowsky, M.J., et al.: Strain tracking reveals the determinants of bacterial engraftment in the human gut following fecal microbiota transplantation. Cell host & microbe 23(2), 229−240 (2018)

[27] Wang, S., Jiang, Y., Li, S.: Pstrain: An iterative microbial strains profiling algorithm for shotgun metagenomic sequencing data. Bioinformatics (2020)

[28] Truong, D.T., Franzosa, E.A., Tickle, T.L., Scholz, M., Weingart, G., Pasolli, E., Tett, A., Huttenhower, C., Segata, N.: Metaphlan2 for enhanced metagenomic taxonomic profiling. Nature methods 12(10), 902−903 (2015)

[29] Bamba, T., Matsuda, H., Endo, M., Fujiyama, Y.: The pathogenic role of bacteroides vulgatus in patients with ulcerative colitis. Journal of gastroenterology 30, 45−47 (1995)

[30] Shiba, T., Aiba, Y., Ishikawa, H., Ushiyama, A., Takagi, A., Mine, T., Koga, Y.: The suppressive effect of bifidobacteria on bacteroides vulgatus, a putative pathogenic microbe in inflammatory bowel disease. Microbiology and immunology 47(6), 371−378 (2003)

[31] Quan, Y., Song, K., Zhang, Y., Zhu, C., Shen, Z., Wu, S., Luo, W., Tan, B., Yang, Z., Wang, X.: Roseburia intestinalis-derived flagellin is a negative regulator of intestinal inflammation. Biochemical and biophysical research communications 501(3), 791−799 (2018)

[32] Shen, Z., Zhu, C., Quan, Y., Yang, J., Yuan, W., Yang, Z., Wu, S., Luo, W., Tan, B., Wang, X.: Insights into roseburia intestinalis which alleviates experimental colitis pathology by inducing anti-inflammatory responses. Journal of gastroenterology and hepatology 33(10), 1751−1760 (2018)

[33] Xiao, M., Shen, Z., Luo, W., Tan, B., Meng, X., Wu, X., Wu, S., Nie, K., Tong, T., Hong, J., et al.: A new colitis therapy strategy via the target colonization of magnetic nanoparticle-internalized roseburia intestinalis. Biomaterials science 7(10), 4174−4185 (2019)

[34] Zhu, C., Song, K., Shen, Z., Quan, Y., Tan, B., Luo, W., Wu, S., Tang, K., Yang, Z., Wang, X.: Roseburia intestinalis inhibits interleukin-17 excretion and promotes regulatory t cells differentiation in colitis. Molecular Medicine Reports 17(6), 7567−7574 (2018)

[35] Samarkos, M., Mastrogianni, E., Kampouropoulou, O.: The role of gut microbiota in clostridium difficile infection. European journal of internal medicine 50, 28−32 (2018)

[36] Song, Y., Garg, S., Girotra, M., Maddox, C., von Rosenvinge, E.C., Dutta, A., Dutta, S., Fricke, W.F.: Microbiota dynamics in patients treated with fecal microbiota transplantation for recurrent clostridium difficile infection. PloS one 8(11), 81330 (2013)

[37] Vaughn, B.P., Vatanen, T., Allegretti, J.R., Bai, A., Xavier, R.J., Korzenik, J., Gevers, D., Ting, A., Robson, S.C., Moss, A.C.: Increased intestinal microbial diversity following fecal microbiota transplant for active crohn’s disease. Inflammatory bowel diseases 22(9), 2182−2190 (2016)

[38] Huang, H.L., Chen, H.T., Luo, Q.L., Xu, H.M., He, J., Li, Y.Q., Zhou, Y.L., Yao, F., Nie, Y.Q., Zhou, Y.J.: Relief of irritable bowel syndrome by fecal microbiota transplantation is associated with changes in diversity and composition of the gut microbiota. Journal of digestive diseases 20(8), 401−408 (2019)

[39] Jia, W., Li, H., Li, S., Chen, L., Li, S.C.: Oviz-bio: a web-based platform for interactive cancer genomics data visualization. Nucleic acids research 48(W1), 415−426 (2020)

[40] Stackebrandt, E., Goebel, B.M.: Taxonomic note: a place for dna-dna reassociation and 16s rrna sequence analysis in the present species definition in bacteriology. International journal of systematic and evolutionary microbiology 44(4), 846−849 (1994)

[41] Tariq, R., Saha, S., Solanky, D., Pardi, D.S., Khanna, S.: Predictors and management of failed fecal microbiota transplantation for recurrent clostridioides difficile infection. Journal of Clinical Gastroenterology 55(6), 542−547 (2021)

[42] Allegretti, J.R., Allegretti, A.S., Phelps, E., Xu, H., Fischer, M., Kassam, Z.: Classifying fecal microbiota transplantation failure: An observational study examining timing and characteristics of fecal microbiota transplantation failures. Clinical gastroenterology and hepatology: the official clinical practice journal of the American Gastroenterological Association 16(11), 1832−1833 (2017)

[43] Kachlíková, M., Sabaka, P., Koščálová, A., Bendžala, M., Dovalová, Z., Stankovič, I.: Comorbid status and the faecal microbial transplantation failure in treatment of recurrent clostridioides difficile infection−pilot prospective observational cohort study. BMC infectious diseases 20(1), 1−9 (2020)

[44] Fischer, M., Kao, D., Mehta, S.R., Martin, T., Dimitry, J., Keshteli, A.H., Cook, G.K., Phelps, E., Sipe, B.W., Xu, H., et al.: Predictors of early failure after fecal microbiota transplantation for the therapy of clostridium difficile infection: a multicenter study. American Journal of Gastroenterology 111(7), 1024−1031 (2016)

[45] Team, R.C., et al.: R: A language and environment for statistical computing (2013)

[46] Kolde, R., Kolde, M.R.: Package ‘pheatmap’. R package 1(7), 790 (2015)

[47] Benson, D.A., Cavanaugh, M., Clark, K., Karsch-Mizrachi, I., Lipman, D.J., Ostell, J., Sayers, E.W.: Genbank. Nucleic acids research 41(D1), 36−42 (2012)

[48] Delcher, A.L., Phillippy, A., Carlton, J., Salzberg, S.L.: Fast algorithms for large-scale genome alignment and comparison. Nucleic acids research 30(11), 2478−2483 (2002)

[49] Price, M.N., Dehal, P.S., Arkin, A.P.: Fasttree 2−approximately maximum-likelihood trees for large alignments. PloS one 5(3), 9490 (2010)

[50] Wickham, H.: Ggplot2: Elegant Graphics for Data Analysis. Springer, ??? (2016). https://ggplot2.tidyverse.org

